# Hepatocyte FGFR2 Mediates the Antifibrotic Effects of FGF10 in Advanced Liver Fibrosis

**DOI:** 10.1101/2025.11.18.689001

**Authors:** Xuanxin Yang, Bingjie Yu, Jiaying Ma, Yawei Yan, Huan Wang, Qingqing Dong, Shuangyan Peng, Qiqi Wu, Tingting Zhang, Panyu Zhang, Zelong Jiang, Chao Lu, Le Li, Xinyi You, Yuandong Xu, Joshua Banda, Zhixiang Mu, Minghua Zheng, Xiaokun Li, Qi Hui, Xiaojie Wang

## Abstract

Fibroblast growth factor 10 (FGF10) supports epithelial repair, but its role in liver fibrogenesis remains uncertain. Here we found that hepatic FGF10 and its receptor FGFR2 decline with progression to advanced fibrosis in patient biopsies and in carbon tetrachloride (CCl4)- and diet-induced mouse models. Restoring hepatic FGF10 with recombinant human FGF10 or adeno-associated virus mediated liver-targeted expression attenuated and partially reversed established advanced bridging fibrosis, reduced inflammation, and decreased hepatocyte apoptosis, and these benefits required hepatocyte FGFR2 because hepatocyte-specific Fgfr2 deletion abolished protection. In primary hepatocytes, FGF10 activated FGFR2-FRS2α, increased inhibitory GSK3β(Ser9) phosphorylation, and suppressed NF-κB activation, reducing TGF-β1 and other cytokines and thereby limiting paracrine hepatic stellate cells activation. The molecular and histologic benefits extended to metabolically primed steatohepatitis, supporting translatability beyond toxicant injury. Mechanistic signaling readouts confirm FGFR2 engagement, while the main text centers on the hepatocyte FGF10-FGFR2 axis as an intrinsic brake on fibrogenesis. Together, these data identify a hepatocyte-centric FGF10-FGFR2 axis as a negative regulator and potential regressor of fibrogenesis, highlighting it as a tractable therapeutic target in advanced liver fibrosis.

## INTRODUCTION

Metabolic dysfunction-associated steatotic liver disease (MASLD) is now the most common chronic liver disorder, affecting approximately 30% of adults worldwide (Stefan Yki-Järvinen & Neuschwander-Tetri, 2025; Wong *et al*, 2023; Younossi *et al*, 2023). Although simple steatosis is often clinically silent, a subset of patients progresses to metabolic dysfunction-associated steatohepatitis (MASH), characterized by inflammation, ballooning injury, and, crucially, progressive fibrosis (Man *et al*, 2023; Ng *et al*, 2023; Simon *et al*, 2023). The rising prevalence of MASLD, coupled with the lack of approved antifibrotic therapies, underscores the urgency of identifying molecular pathways that determine fibrotic progression (Harrison *et al*, 2024; Schwabe *et al*, 2025; Steinberg Carpentier & Wang, 2025). Fibrosis stage is the strongest histologic predictor of liver-related and overall mortality in MASLD, making pathways that modulate fibrotic burden particularly attractive for therapeutic exploration.

Liver fibrosis is initiated by hepatocyte injury and sustained by paracrine activation of hepatic stellate cells (HSCs). Injured hepatocytes release inflammatory mediators and undergo apoptosis, processes regulated by intracellular signaling pathways such as nuclear factor kappa B (NF-κB) and glycogen synthase kinase 3β (GSK3β) (Du *et al*, 2023; Taru *et al*, 2024; Wang *et al*, 2025). This dysfunctional state increases secretion of transforming growth factor-β1 (TGF-β1) and other cytokines (Gaul *et al*, 2021; Hammerich & Tacke, 2023; Taru *et al*., 2024), which drive transdifferentiation of HSCs into myofibroblast-like cells that produce extracellular matrix (ECM) (Bourebaba & Marycz, 2021; Payen *et al*, 2021; Yang *et al*, 2021b). Therefore, strategies that preserve hepatocyte viability and suppress inflammation are expected to mitigate fibrogenesis by re-establishing hepatocyte control over HSC activation and the fibrotic niche.

Fibroblast growth factors (FGFs) orchestrate development, tissue homeostasis, and repair in multiple organs. In the liver, endocrine FGFs such as FGF19 and FGF21 modulate bile acid and metabolic homeostasis and are being evaluated as therapeutic candidates in MASLD (Alvarez-Sola *et al*, 2017; Li *et al*, 2024; Tan *et al*, 2023). In contrast, the contributions of paracrine FGFs to chronic liver injury are less defined. Fibroblast growth factor 10 (FGF10), a prototypical paracrine FGF, primarily signals through epithelial FGFR2 (Berg *et al*, 2007; Itoh & Ohta, 2014; Yang *et al*, 2021a), and supports tissue regeneration (Yang *et al*, 2020; Zhou *et al*, 2022), yet its function in chronic liver fibrogenesis and the requirement for hepatocyte FGFR2 remain insufficiently defined.

We found that hepatic FGF10 and FGFR2 are reduced in MASLD, particularly in advanced fibrosis, consistent with reports that FGF10 limits apoptosis and dampens inflammatory signaling in other tissues (Chen *et al*, 2017; Dong *et al*, 2019), suggesting that this axis may serve as an intrinsic epithelial brake on injury-induced inflammation. We hypothesized that chronic injury downregulates the FGF10-FGFR2 axis within hepatocytes, thereby removing a cytoprotective brake on inflammatory and apoptotic signaling and increasing paracrine activation of hepatic stellate cells. We further posited that augmenting FGF10 would suppress inflammation and apoptosis in a hepatocyte FGFR2-dependent manner, leading to inhibition and partial reversal of established fibrosis even at the stage of bridging fibrosis. Thus, we investigated the hepatocyte FGF10-FGFR2 axis as a hepatocyte-centered therapeutic entry point to re-establish epithelial control of the fibrotic niche in advanced liver fibrosis.

## RESULTS

### Hepatic FGF10 and FGFR2 decrease with fibrosis in patients

We profiled hepatic FGF10 across fibrosis stages by integrating transcript and protein analyses of fresh and formalin-fixed, paraffin-embedded (FFPE) human liver biopsies. In fresh liver biopsies (n = 27), *FGF10* mRNA levels inversely correlated with fibrosis stage (Fig. 1A and 1B), whereas FGF2 and FGF21 transcripts showed no linear relationship with fibrosis score (Appendix Fig. S1), indicating specificity of the FGF10-fibrosis score association. Quantitative immunohistochemistry on FFPE samples detected FGF10 protein at fibrosis stages (FS) 0 to 3 and revealed a stepwise reduction in staining intensity with increasing fibrosis (Fig. 1C and 1D; Appendix Fig. S2). Histopathology confirmed progressive collagen deposition from FS0 through FS3, culminating in bridging fibrosis linking central-central and central-portal regions at FS3, a pattern characteristic of advanced liver fibrosis (Appendix Fig. S3). FGFR2 expression also decreased with advancing fibrosis stage (FS3 vs. FS0) in human biopsies (see Fig. 1E and 1F), and these findings aligned with the ligand decline and supported a coordinated reduction of the FGF10-FGFR2 axis that is positioned to weaken hepatocyte-centered control of fibrogenic remodeling.

**Figure 1.**
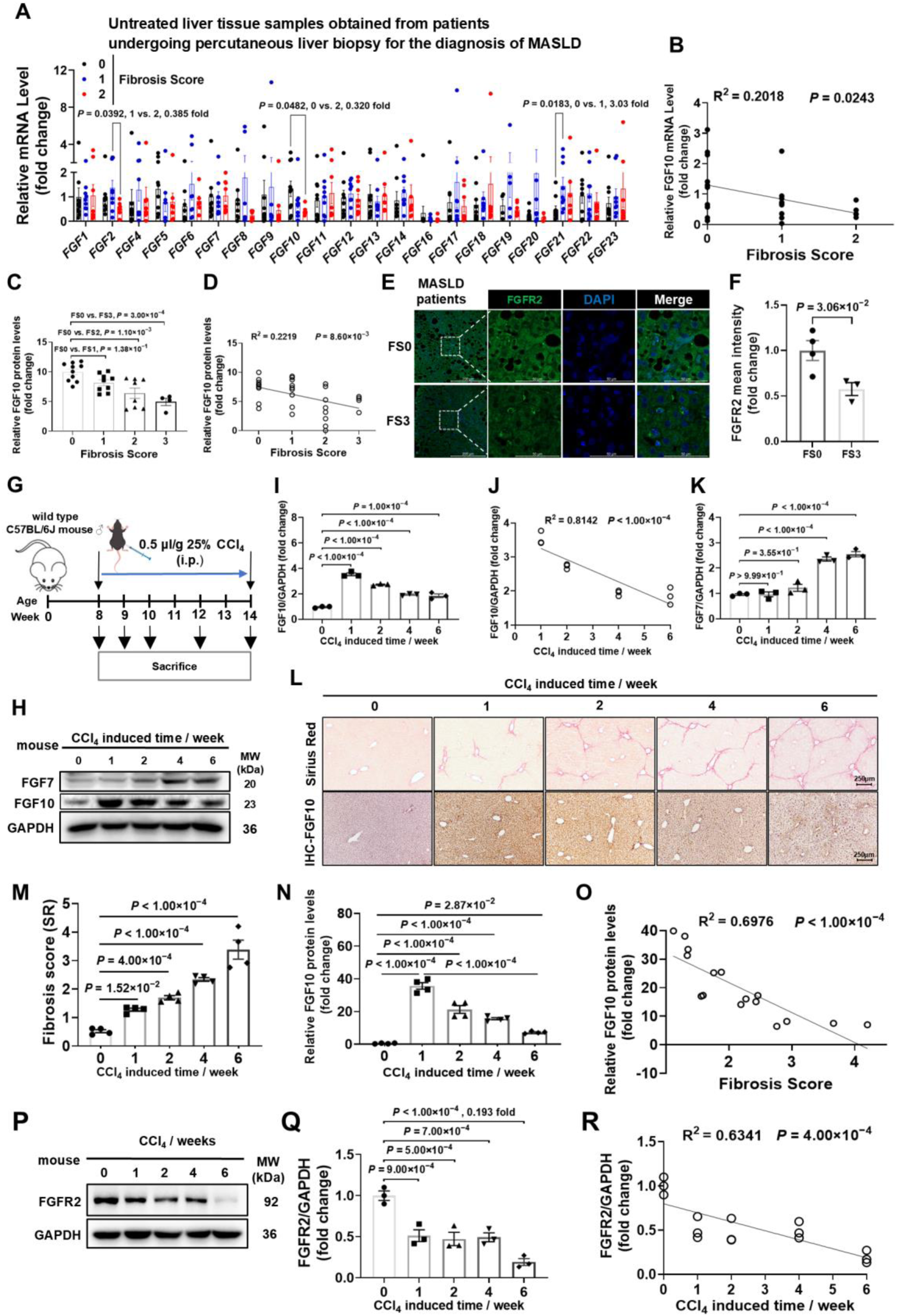
Coordinated decreases of hepatic FGF10 and FGFR2 with fibrosis in patients and in CCl₄ -injured mice. (A) Hepatic FGF transcripts in MASLD biopsies by FS (FS0, n = 10; FS1, n = 9; FS2, n = 8). (B) Linear regression of FGF10 mRNA vs FS (per patient; n = 4-10/group across panels B-D). (C-D) FGF10 IHC across FS0-FS3 with quantification (C) and regression vs FS (D). (E–F) FGFR2 IF in the same cohort across FS0 versus FS3 with quantification. (G) CCl_4_ protocol schematic. (H-K) Time-course immunoblot of hepatic FGF10 and FGF7 over 0-6 weeks; quantifications for FGF10 (I) and FGF7 (K); (J) regression of FGF10 vs CCl_4_ weeks. (L) Representative SR and FGF10 IHC across time; scale bar, 250 μm. (M-N) Quantification of SR% (M) and FGF10 IHC (N) (n = 4/group). (O) Regression of FGF10 protein vs fibrosis score. (P-R) Hepatic FGFR2 protein time course and quantification in the same CCl_4_ series. Stats: unpaired t-test (F), one-way ANOVA (A, C, I, K, M, N, Q).

We next reanalyzed the human liver RNA-seq dataset (GSE246221) and restricted samples to NAS = 5-6. FGFR2 transcripts declined with increasing FS from 1 to 3, whereas FGFR1 showed no significant trend (Appendix Fig. S4). This external validation supports a fibrosis-dependent reduction of FGFR2 under matched disease activity and reinforces attenuation of the epithelial FGF10-FGFR2 axis during progression.

### Hepatic FGF10 and FGFR2 decline in murine models of fibrosis

Given the metabolic context, we first examined FGF10 in high-fat diet (HFD)-fed models. Mice fed an HFD for 5, 16, or 20 weeks exhibited modest reductions in hepatic FGF10 compared with chow-fed controls, without a clear time-dependent trend (Appendix Fig. S5), consistent with the lack of apparent FS3-like bridging patterns (Appendix Fig. S6). We then induced liver fibrosis using a 6-week carbon tetrachloride (CCl_4_) protocol (schema in Fig. 1G). In this model, which develops FS3-like bridging fibrosis, hepatic FGF10 protein was transiently induced at 1 week and then progressively declined from 1 to 6 weeks, resulting in an overall reduction at the stage of bridging fibrosis (Fig. 1H-1J). By contrast, the related family member FGF7 was induced and remained elevated over the same interval (Fig. 1H and 1K; Appendix Fig. S7). Fibrosis severity increased with induction time, and FGF10 abundance showed a strong inverse correlation with histologic stage (Fig. 1L-1O). In parallel, hepatic FGFR2 protein progressively declined over the same CCl_4_ time course (Fig. 1P-1R), further indicating loss of the receptor arm of this pathway during fibrogenesis. In the same series, FGFR1 exhibited only a modest decrease that plateaued at later time points (Appendix Fig. S8). Together, these murine models recapitulate the human pattern of coordinated FGF10-FGFR2 downregulation with fibrosis progression and support the concept that chronic injury erodes an endogenous FGF10-FGFR2 brake in hepatocytes.

### FGF10 localizes predominantly to hepatic stellate cells

To define the cellular source of FGF10 during fibrosis, reanalysis of the single-cell RNA-seq dataset GSE212837 (total of 128,851 cells) likewise mapped FGF10 transcripts mainly to the HSC cluster (Fig. 2A-2B). Stage stratification (Control; MASH FS1-FS3) showed stage-dependent increases of FGF10 transcripts within HSCs (Fig. 2C). To assess this at the tissue level, we mapped its localization in human FS3 biopsies and CCl_4_-injured mouse livers. Immunofluorescence demonstrated predominant colocalization of FGF10 with desmin- and α-smooth muscle actin (α-SMA)-positive HSCs (Fig. 2D). This compartment- and stage-specific pattern provides a framework to reconcile transient FGF10 induction in early injury with loss of the broader FGF10-FGFR2 axis in advanced disease and suggests that epithelial compartments, including hepatocytes, progressively lose access to this paracrine signal.

**Figure 2.**
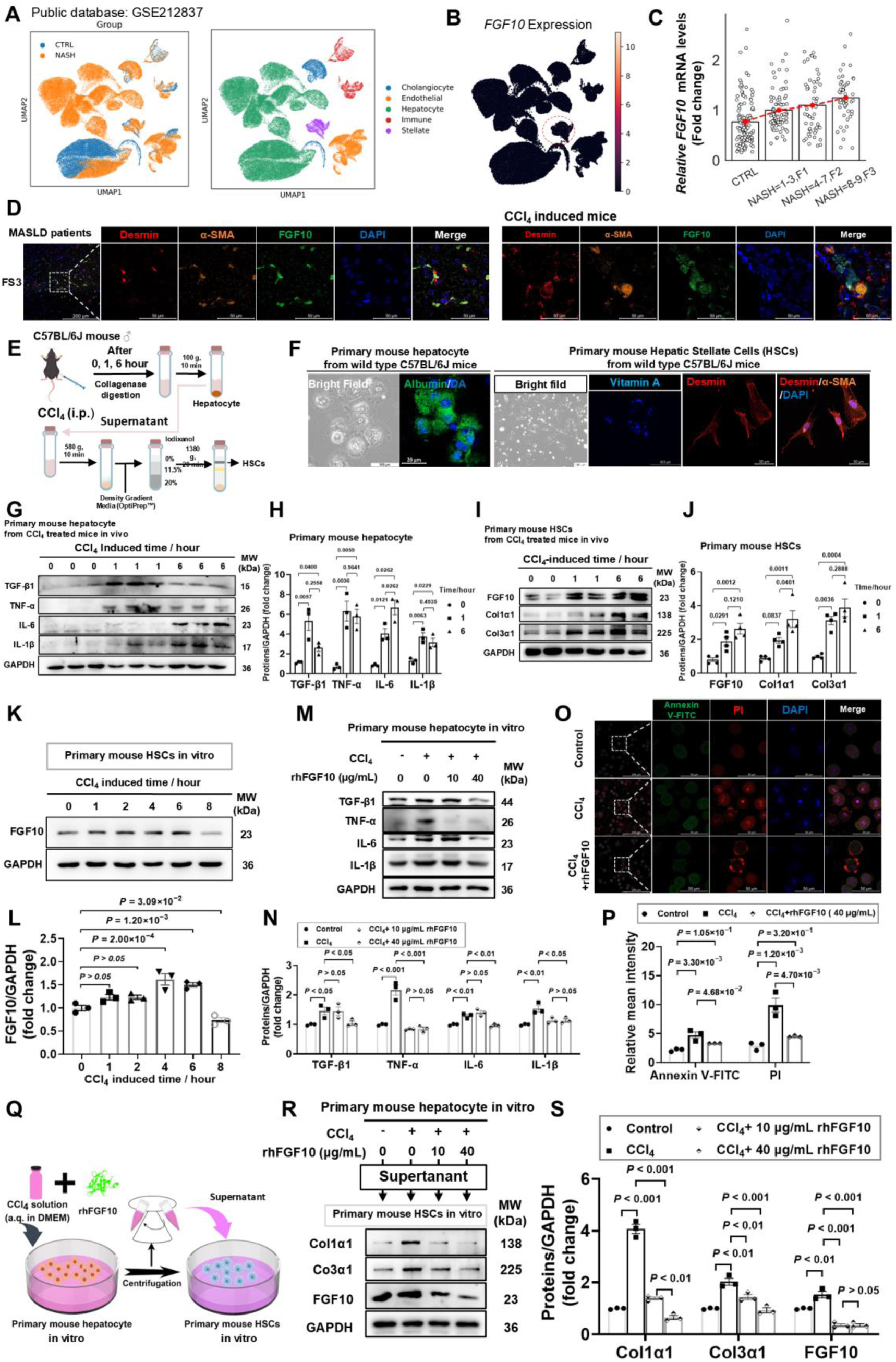
FGF10 reduces hepatocyte inflammatory output and apoptosis and limits paracrine HSC activation in vitro. (A-C) Human liver scRNA-seq (GSE212837): UMAP cell types (A), FGF10 feature map enriched in HSCs (B), and stage-stratified FGF10 in HSCs (C). (D) IF in human FS3 and 6-week CCl_4_ mouse liver showing FGF10 colocalization with desmin/α-SMA in HSCs; scale bars, 200 μm (overview) and 50 μm (channels). (E) Workflow for isolating primary cells from CCl₄ -treated C57BL/6J mouse liver. (F) Primary mouse hepatocytes (ALB+) and HSCs (vitamin A autofluorescence; DESMIN/α-SMA+). (G-H) Hepatocytes collected 0-6 h after CCl_4_ show rising TGF-β1, TNF-α, IL-6, and IL-1β by immunoblot. Quantification is normalized to GAPDH and baseline. (I-J) HSCs from the same livers show altered FGF10 and increased Col1a1 and Col3a1. Quantification is normalized to GAPDH and baseline. (K-L) FGF10 protein dynamics in HSCs after CCl_4_ (0-8 hour) with quantification. (M-N) In CCl_4_-injured hepatocytes, rhFGF10 (0, 10, 40 μg/mL, 24 h) lowers TNF-α, IL-6, IL-1β in a dose-dependent manner. (O-P) Annexin V/PI readouts show reduced apoptosis by rhFGF10; scale bar, 50 μm. (Q) Indirect co-culture schematic in vitro. (R-S) HSCs exposed to hepatocyte-conditioned media (+/-rhFGF10) show reduced schematic. Stats: one-way ANOVA (H, J, L, N, P, S).

### FGF10 limits hepatocyte inflammatory signaling and apoptosis and indirectly restrains stellate-cell activation

We next tested whether FGF10 modulates hepatocyte-HSC communication using primary cells. As shown in the isolation workflow (Fig. 2E), livers were collagenase-perfused, hepatocytes were recovered by low-speed sedimentation, and HSCs were enriched from the supernatant by density gradient, enabling paired sampling from the same injured livers (Fig. 2E). Primary mouse hepatocytes and HSCs were isolated from 8-week-old C57BL/6J mice and validated by lineage-specific markers (Fig. 2F). Within hepatocytes, acute CCl_4_ exposure in vivo (0, 1 and 6 h) rapidly increased inflammatory output, with TGF-β1, TNF-α, IL-6, and IL-1β rising over time by immunoblot (Fig. 2G and 2H). In the matched HSC fraction, FGF10 protein and the fibrosis markers Col1a1 and Col3a1 increased (Fig. 2I and 2J), indicating an early fibrogenic shift accompanied by FGF10 induction. In vitro, CCl_4_ similarly increased TGF-β1 in hepatocytes and altered FGF10 levels in HSCs (Appendix Fig. S9). Time-course immunoblotting in primary HSCs showed a biphasic response: FGF10 rose from 2 to 6 h, peaking at 4 and 6 h, and then declined sharply by 8 h (Fig. 2K and 2L).

Because receptor-binding determinants are conserved and recombinant human FGF10 (rhFGF10) effectively engages the murine receptor, we used rhFGF10 for in vitro stimulation. In CCl_4_-injured hepatocytes, rhFGF10 at 10 and 40 μg/mL reduced TGF-β1, TNF-α, IL-6, and IL-1β in a concentration-dependent manner, with maximal suppression at 40 μg/mL (Fig. 2M and 2N). Annexin V-FITC/propidium iodide staining demonstrated reduced apoptosis with rhFGF10 treatment (Fig. 2O and 2P).

To assess paracrine effects on HSCs, conditioned medium from hepatocytes treated with CCl_4_ ± rhFGF10 was applied to primary HSCs (Fig. 2Q). Relative to medium from CCl_4_-only hepatocytes, rhFGF10-conditioned medium decreased Col1a1 and Col3a1 expression and reduced FGF10 levels in HSCs (Fig. 2R and 2S), consistent with attenuated HSC activation. While some indicators exhibited modest variability across doses, the overall direction of effect was consistent and compatible with biological heterogeneity. Taken together, these data support a model in which FGF10 signaling within hepatocytes dampens cytokine output and apoptosis and thereby imposes a hepatocyte-centered paracrine brake on HSC activation.

### Hepatic FGF10 overexpression downregulates profibrotic signaling and ECM programs in CCl_4_-induced liver fibrosis

To test the functional impact of hepatic FGF10, we achieved liver-directed overexpression using an AAV2/8 vector encoding *Fgf10* (AAV-*Fgf10*)and AAV-GFP as the control (Li *et al*, 2021). Intravenous delivery produced a liver-restricted fluorescent signal from day 7 to day 28, and ex vivo imaging confirmed predominant hepatic transduction without detectable signal in non-hepatic organs (Fig. 3A). Consistently, hepatic FGF10 protein and *Fgf10* mRNA were increased in AAV-*Fgf10* mice (Appendix Fig. S10A-S10C). After a 3-week expression window, mice underwent a 6-week CCl_4_ injury protocol (Fig. 3B). RNA sequencing (RNA-seq) of liver tissue showed clear group separation by principal component analysis, with high within-group concordance (Fig. 3C and 3D). Using DESeq2 with BH correction, we identified 1,644 DEGs (FDR < 0.05; |log2FC| ≥ 0.58) between AAV-*Fgf10* and AAV-GFP livers (695 up, 949 down; Fig. 3E).

**Figure 3.**
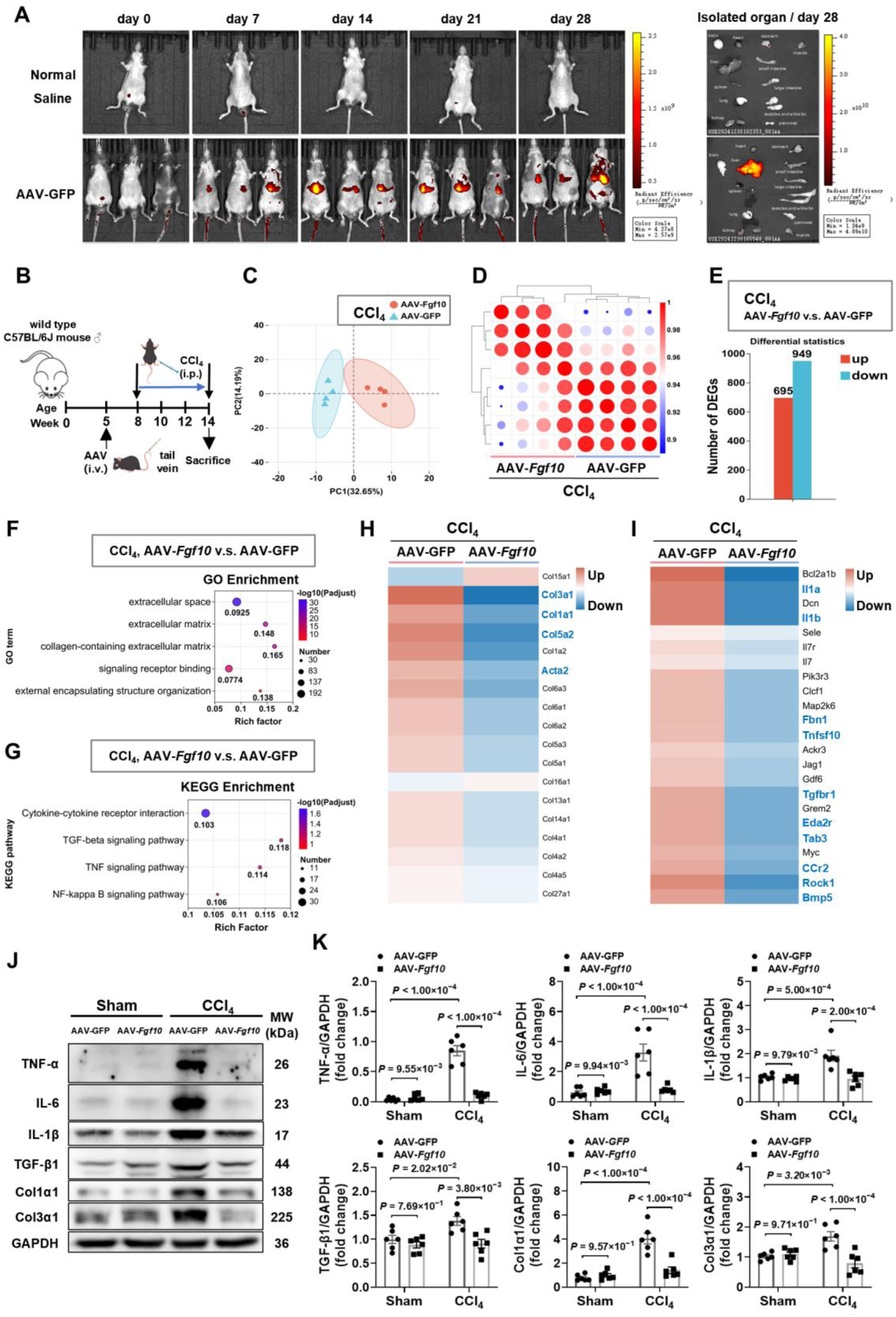
FGF10 reprograms inflammatory and ECM programs in CCl_4_-injured mouse liver. (A) In vivo/ex vivo AAV-GFP signals over 28 days post-i.v. injection. (B) Study schema: AAV (week 5) and CCl_4_ (weeks 8-14). (C-E) PCA separation, replicate correlation, and DEG counts (AAV-*Fgf10* vs AAV-GFP; n = 4/group). (F-G) GO and KEGG enrichment highlight ECM and cytokine, TGF-β, TNF pathways. (H-I) Heatmaps of collagen, Acta2 and cytokine, TGF-β, TNF modules. (J-K) Immunoblots and quantification of TNF-α, IL-6, IL-1β, TGF-β1, Col1α1, Col3α1 (sham vs CCl_4_; AAV-GFP vs AAV-*Fgf10*; n = 6/group). Stats: two-way ANOVA (K); RNA-seq DEGs by FDR < 0.05, |log2FC| ≥ 0.58.

Gene Ontology enrichment analysis highlighted collagen-containing ECM and external encapsulating structure terms (Fig. 3F), and hierarchical clustering showed downregulation of fibrogenic markers, including *Col3a1*, *Col1a1*, *Col5a2*, and *Acta2* (Fig. 3H). KEGG analysis indicated modulation of cytokine-receptor interaction and stress-response pathways linked to TGF-β, TNF, and NF-κB (Fig. 3G); within these axes, representative pro-apoptotic (e.g., *Tnfsf10*, *Rock1*, *Eda2r*) and pro-inflammatory transcripts (e.g., *Il1a*, *Il1b*, *Ccr2*, *Tab3*) were reduced in AAV-*Fgf10* livers (Fig. 3I). Immunoblotting validated lower ECM protein abundance (Col1α1, Col3α1) and reduced TGF-β1, TNF-α, IL-6, and IL-1β (Fig. 3J and 3K), consistent with attenuated TGF-β/TNF/NF-κB signaling in vivo. Thus, hepatocyte-directed FGF10 overexpression broadly reprograms profibrotic and inflammatory transcriptional programs in the fibrotic liver, consistent with restoration of an epithelial antifibrotic circuit.

### Liver-restricted FGF10 attenuates bridging fibrosis in CCl_4_-injured mice

We next assessed whether hepatocyte-directed FGF10 modifies histological injury and fibrogenesis. Relative to AAV-GFP, AAV-*Fgf10* treatment improved hepatic architecture and reduced ballooning on H&E staining (Fig. 4A). Sirius Red staining and quantitative scoring showed less collagen deposition and lower fibrosis stage (Fig. 4B and 4C). Immunohistochemistry revealed fewer α-SMA- and TGF-β1-positive cells, consistent with reduced HSC activation (Fig. 4D-4F), and diminished macrophage infiltration by F4/80 staining (Fig. 4G and 4H). Systemic indices, including body weight, liver weight, and liver-to-body-weight ratio, were comparable at the endpoint (Fig. 4I and 4J), indicating that FGF10 overexpression did not induce hepatomegaly or alter gross growth parameters. Serum alanine aminotransferase (ALT) and aspartate aminotransferase (AST) remained elevated in both groups without significant between-group differences (Fig. 4K), suggesting that biochemical normalization may lag behind histologic improvement under these conditions. Relative to time-matched CCl_4_ controls, collagen burden and fibrosis stage were reduced in AAV-*Fgf10* mice, indicating partial reversal rather than only slowed progression and supporting the concept that structural repair can precede biochemical normalization in chronic injury. These findings link restoration of the hepatocyte FGF10-FGFR2 axis to macroscopic regression of bridging fibrosis, consistent with hepatocyte-centered control of matrix remodeling.

**Figure 4.**
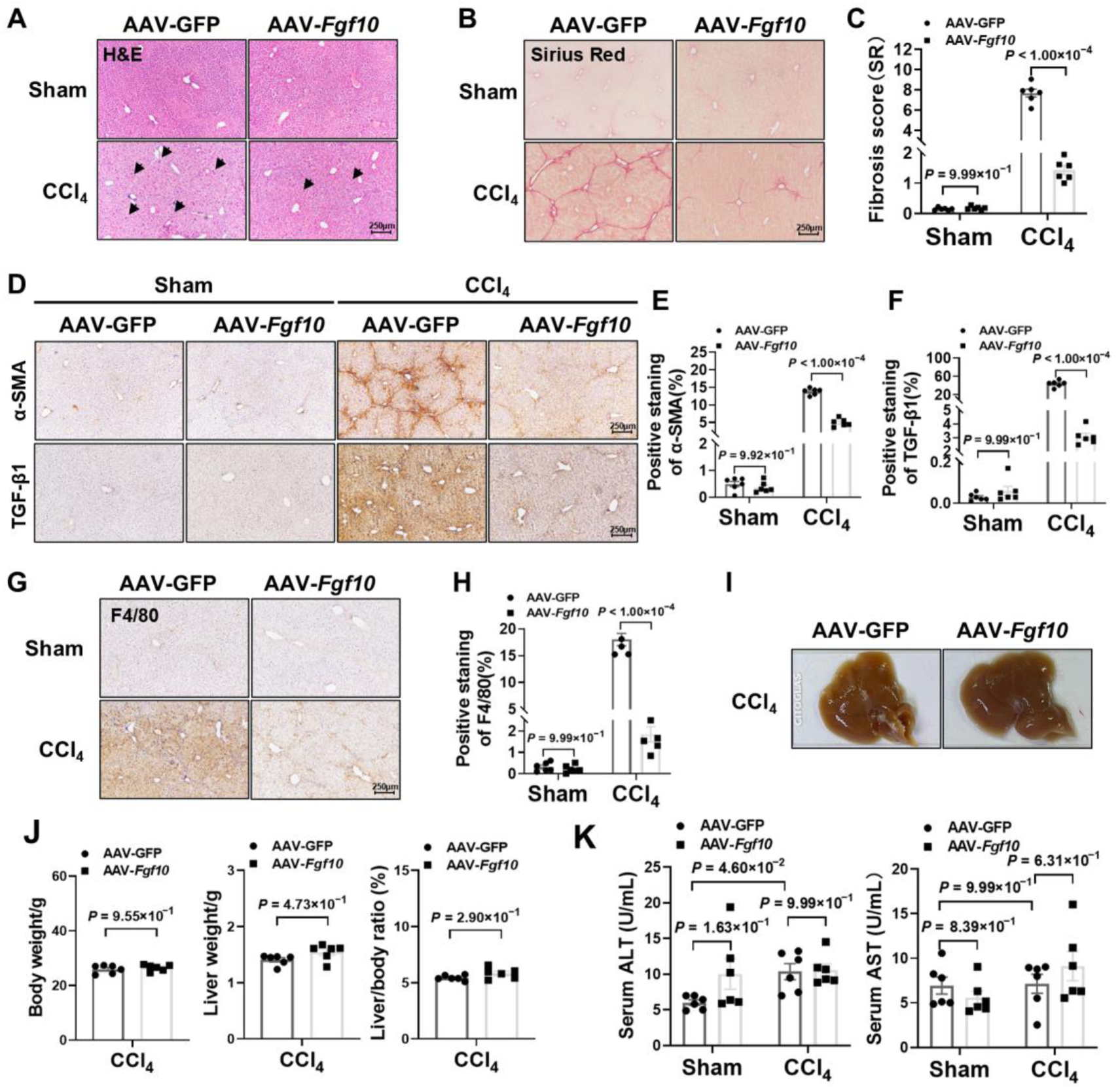
Liver-restricted FGF10 mitigates CCl_4_-induced injury and fibrosis. (A) H&E (ballooning indicated); scale bar, 250 μm (n = 6/group). (B-C) SR and morphometry. (D-H) IHC and quantification of α-SMA, TGF-β1, and F4/80. (I-J) Gross liver images and body/liver indices. (K) Serum ALT/AST. Stats: unpaired t-test (C, E, F, H, J); two-way ANOVA (K).

### Liver-restricted FGF10 mitigates fibrosis and steatohepatitis in a high-fat diet plus CCl_4_ model

To evaluate efficacy in a metabolic-toxicant context, mice received 20 weeks of high-fat diet followed by 8 weeks of CCl_4_, and were transduced with AAV-*Fgf10* for hepatocyte-directed FGF10 expression (Appendix Fig. S11A). AAV-*Fgf10* reduced steatohepatitis and collagen deposition with less bridging fibrosis on histology (Appendix Fig. S11B and S11C). These changes were accompanied by downregulation of lipogenic and cholesterogenic genes, including *Fasn* and *Srebf2*, with a downward trend in *Mvk* (Appendix Fig. S11D). Macroscopic liver appearance improved (Appendix Fig. S11E). Despite no difference in body weight between groups, AAV-*Fgf10* mice displayed higher liver weight and liver-to-body-weight ratio at endpoint (Appendix Fig. S11F), a pattern consistent with restoration of hepatic mass during injury resolution rather than pathological enlargement. Serum AST and ALT were significantly lower with AAV-*Fgf10* (Appendix Fig. S11G). At the protein level, hepatic FGF10 overexpression suppressed profibrotic and inflammatory mediators, including Col1a1, α-SMA, fibronectin, TNF-α, and IL-1β, and reduced the pro-apoptotic marker Bax (Appendix Fig. S11H and S11I), aligning molecular, biochemical, and histologic endpoints. Together with the CCl_4_-only model, these data indicate that reinforcing hepatocyte FGF10-FGFR2 signaling preserves antifibrotic efficacy and hepatocyte survival in metabolically primed steatohepatitis, extending the concept of hepatocyte-centered control of fibrosis beyond pure toxicant injury.

### FGF10 partially restores FGFR2 expression in liver fibrosis

To evaluate receptor-level remodeling, we profiled hepatic transcripts in CCl_4_-treated mice expressing either AAV-*Fgf10* or AAV-GFP (Fig. 5A). As expected, *Fgf10* was among the most upregulated transcripts and was accompanied by increased *Fgfr2* expression. Several receptor tyrosine kinases linked to fibrogenesis (*Pdgfra*, *Fgfr3*, *Erbb4*, *Epha3*, *Epha7*, *Ntrk2*, *Mup12*) were reduced, whereas *Erbb3* was elevated, indicating a selective strengthening of the FGFR2 axis with FGF10 overexpression. Baseline ligand and receptor decreases in liver fibrosis are presented in Fig. 1. Here, we therefore focus on receptor remodeling under FGF10 augmentation. Together, these data show selective reinforcement and partial restoration of the FGFR2 axis with FGF10 overexpression.

**Figure 5.**
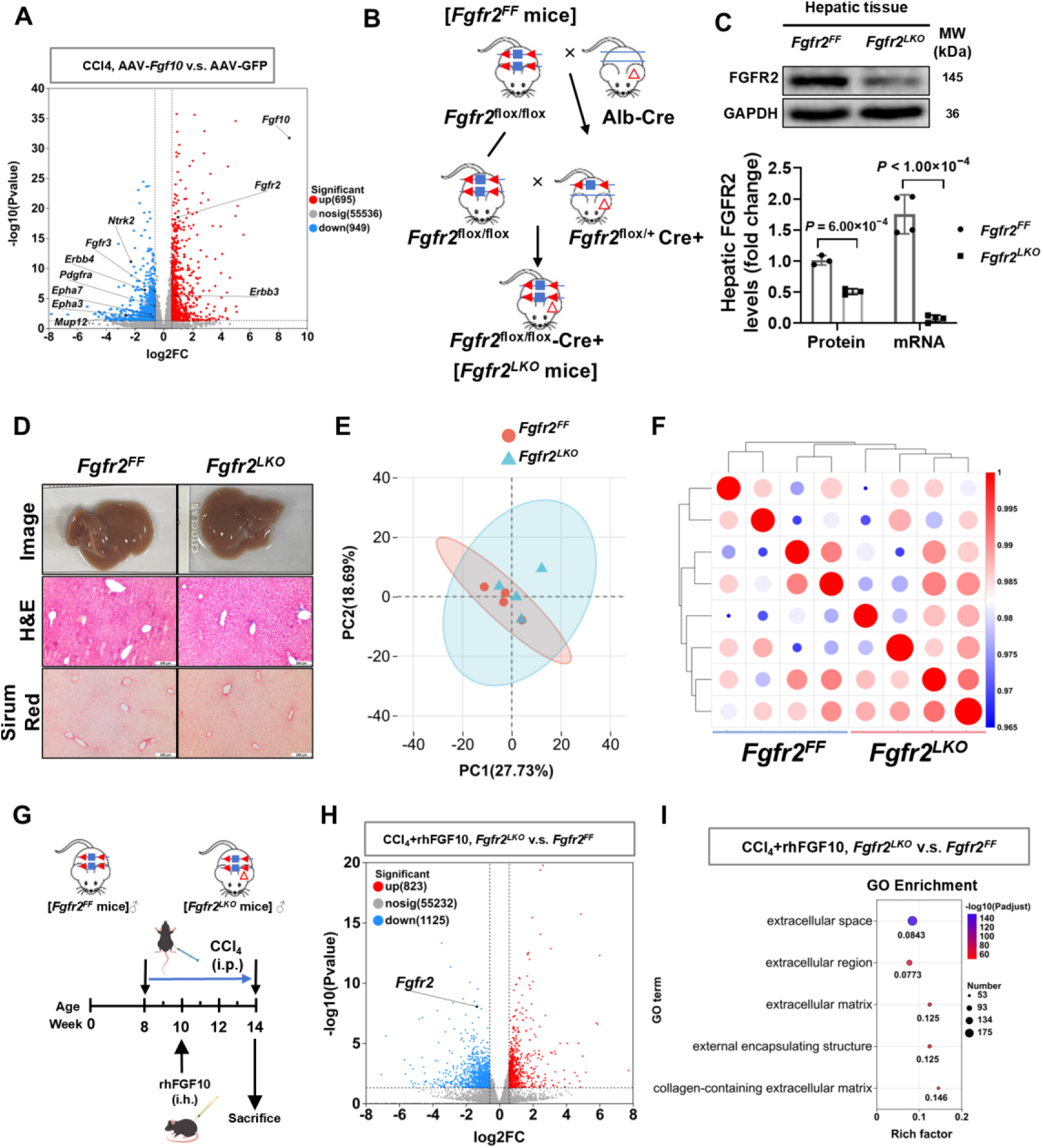
FGFR2 axis remodeling under FGF10 augmentation and baseline characterization of hepatocyte-specific Fgfr2 deletion. (A) Volcano plot of DEGs (AAV-*Fgf10* vs AAV-GFP; n = 4/group). (B) Breeding scheme for *Fgfr2^LKO^* and *Fgfr2^FF^*. (C) Validation of hepatic FGFR2 loss at mRNA and protein levels. (D-F) Basal histology and transcriptomes in *Fgfr2^LKO^*vs *Fgfr2^FF^* show no spontaneous differences. (G) Experimental schema for CCl_4_ + rhFGF10 comparison in *Fgfr2^FF^*vs *Fgfr2^LKO^*. (H) RNA-seq volcano plot for CCl_4_ + rhFGF10: *Fgfr2^LKO^* vs *Fgfr2^FF^*. (I) GO enrichment highlights ECM and collagen-containing structures. Stats: unpaired t-test (C); RNA-seq DEGs by FDR < 0.05.

### Hepatocyte FGFR2 is required for FGF10 efficacy and reshapes fibrosis-related transcriptomes

To test receptor dependence in vivo, we generated hepatocyte-specific Fgfr2 knockout mice (*Fgfr2^LKO^*) by crossing *Fgfr2^flox/flox^* (*Fgfr2^FF^*) with Alb-Cre (Fig. 5B). Efficient deletion was confirmed by genotyping and by reduced hepatic FGFR2 protein and mRNA levels (Appendix Fig. S12 and Fig. 5C). Under basal conditions, gross morphology, H&E, Sirius Red staining, and transcriptomic profiles were comparable between *Fgfr2^LKO^* and *Fgfr2^FF^* (Fig. 5D-5F), indicating that hepatocyte FGFR2 loss does not substantially disrupt liver homeostasis in the absence of injury.

To avoid AAV-related confounders and to standardize exposure across genotypes, we administered rhFGF10 directly in the CCl_4_ model. Daily rhFGF10 administration dose-dependently reduced collagen deposition in CCl_4_-injured mouse livers, with maximal efficacy at 5 mg/kg (Appendix Fig. S13A-S13C), and the administered dose inversely correlated with Sirius Red-positive area (Appendix Fig. S13D). Guided by these data, we compared CCl_4_ + rhFGF10 (5 mg/kg) responses between *Fgfr2^LKO^* and *Fgfr2^FF^* (Fig. 5G). RNA-seq revealed a markedly altered transcriptional response in the absence of hepatocyte FGFR2, with 1,125 genes downregulated and 823 upregulated in *Fgfr2^LKO^*versus *Fgfr2^FF^* (Fig. 5H). Gene Ontology enrichment analysis highlighted extracellular space, ECM, and collagen-containing structures (Fig. 5I), supporting hepatocyte FGFR2 as a key conduit through which FGF10 reprograms fibrotic gene networks. These findings indicate that hepatocyte FGFR2 is required for FGF10 to establish a hepatocyte-driven transcriptional state that disfavors ECM accumulation and fibrogenic remodeling.

### rhFGF10 reduces hepatic injury and fibrogenesis in an FGFR2-dependent manner

We next assessed whether the protective effects of rhFGF10 required hepatocyte FGFR2 during CCl_4_-induced fibrosis. Liver weight and liver-to-body-weight ratio did not differ between *Fgfr2^LKO^* and *Fgfr2^FF^*at the endpoint under rhFGF10 treatment (Fig. 6A and 6B). Histologically, *Fgfr2^FF^*mice treated with rhFGF10 showed reduced hepatocyte injury on H&E and lower collagen deposition and fibrosis scores on Sirius Red (Fig. 6C and 6D), whereas *Fgfr2^LKO^*mice did not show these benefits (Fig. 6C and 6D). Concordantly, rhFGF10 decreased α-SMA and TGF-β1 and diminished F4/80-positive macrophage infiltration in *Fgfr2^FF^*, but not in *Fgfr2^LKO^*(Fig. 6E and 6F). TUNEL staining showed reduced hepatocyte apoptosis with rhFGF10 in *Fgfr2^FF^*but not in *Fgfr2^LKO^* (Fig. 6G and 6H). At the protein level, rhFGF10 lowered fibrosis markers (Col1α1, Col3α1, α-SMA) and pro-inflammatory cytokines (TGF-β1, IL-6, IL-1β) in *Fgfr2^FF^*, whereas these suppressive effects were diminished or absent in *Fgfr2^LKO^* (Fig. 6I and 6J). These findings establish the dependence of rhFGF10’s antifibrotic and anti-inflammatory activities on FGFR2 in vivo. In combination with the transcriptional data, they support a model in which hepatocyte FGFR2 is the essential epithelial receptor that allows FGF10 to exert hepatocyte-centered control over inflammatory and fibrotic remodeling.

**Figure 6.**
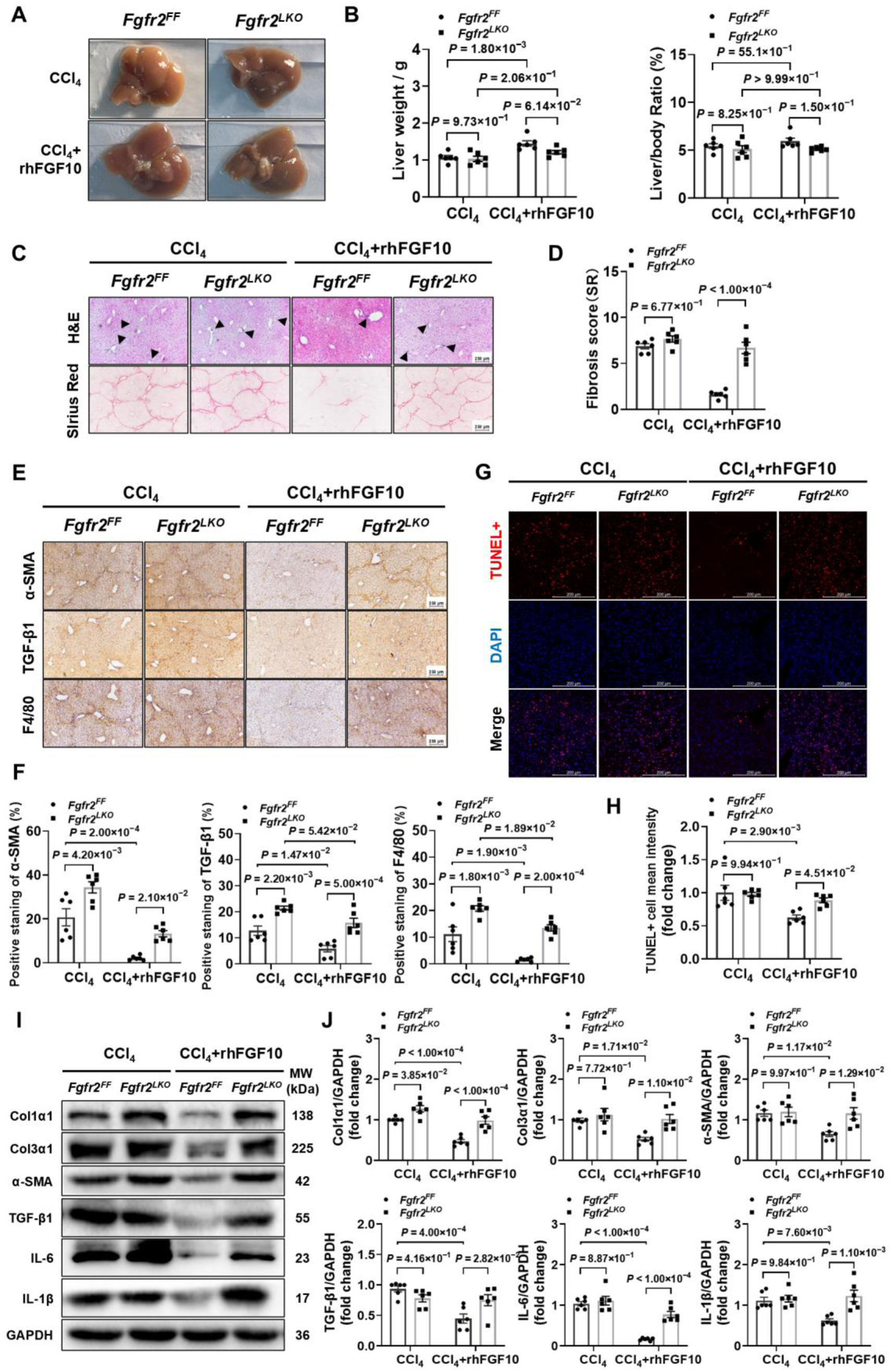
Hepatocyte FGFR2 is required for the antifibrotic, anti-inflammatory, and anti-apoptotic actions of FGF10 in CCl_4_-injured mouse liver. (A) Representative livers (n = 4/group). (B) Liver and liver/body mass indices (n = 6/group). (C-D) H&E/SR and SR morphometry; scale bar, 250 μm. (E-F) IHC/quantification of α-SMA, TGF-β1, F4/80 (n = 6/group). (G-H) TUNEL images/quantification (n = 6/group); scale bar, 50 μm. (I-J) Immunoblots/quantification for Col1α1, Col3α1, α-SMA, TGF-β1, IL-6, IL-1β (n = 6/group). Stats: two-way ANOVA (B, F, H, J); unpaired t-test (D).

### Hepatocyte FGFR2 mediates FGF10-driven FRS2α-GSK3β activation and NF-κB inhibition

To delineate downstream mechanisms, we analyzed key effectors in primary cells after CCl_4_ exposure. In primary mouse hepatocytes, CCl_4_ reduced phosphorylation of FGFR2 (Ser782), with minimal changes in HSCs (Appendix Fig. S14A and S14B), indicating a hepatocyte-centric effect. rhFGF10 restored FGFR2 phosphorylation in CCl_4_-treated hepatocytes to near control (siNC) levels (Appendix Fig. S14C and S14D). Silencing Fgfr2 (si*Fgfr2*) blunted rhFGF10-induced phosphorylation of FRS2α (Tyr196) and GSK3β (Ser9) (Appendix Fig. S14C-S14F), identifying FRS2α-GSK3β as FGFR2-dependent transducers of FGF10 signaling in injured hepatocytes. Because inflammatory signaling fuels fibrogenesis, we assessed NF-κB activity. rhFGF10 reduced p65 phosphorylation and stabilized IκBα (Appendix Fig. S15A and S15B), and these anti-inflammatory effects were largely lost after FGFR2 knockdown in hepatocytes (Appendix Fig. S15C and S15D). Together, these data position hepatocyte FGFR2 upstream of FRS2α-GSK3β activation and NF-κB inhibition in the FGF10 response(Appendix Fig. S15E). In the context of our in vivo findings, they support a mechanistic model in which re-engagement of the hepatocyte FGF10-FGFR2-FRS2α-GSK3β axis restores an epithelial pro-survival, anti-inflammatory program that reasserts hepatocyte-centered control of the fibrotic niche.

## DISCUSSION

This study defines a hepatocyte-centric FGF10-FGFR2 pathway that restrains inflammatory signaling, preserves hepatocyte viability, and limits and can partially reverse fibrotic remodeling. Across patient biopsies and complementary murine models, hepatic FGF10 and FGFR2 declines with advancing stage. Increasing FGF10, by liver-directed gene delivery or recombinant protein, reduced hepatocyte apoptosis, inflammatory infiltration, and ECM deposition. Conversely, hepatocyte-specific loss of Fgfr2 abrogated these benefits, establishing FGFR2 as the required receptor for FGF10-mediated protection during chronic injury. In addition to slowing progression, histologic endpoints demonstrated regression relative to time-matched controls, which supports a partial reversal component under FGF10 augmentation and elevates the FGF10-FGFR2 axis from a protective pathway to a hepatocyte-centered reversal strategy.

Mechanistically, FGF10 engages FGFR2/FRS2α signaling in hepatocytes, increases GSK3β (Ser9) inhibitory phosphorylation, and suppresses NF-κB activity. This cascade maintains epithelial integrity, lowers cytokine output, including TGF-β1, and indirectly limits HSC activation and collagen production. The data align with established roles of GSK3β and NF-κB in hepatocyte stress responses and inflammatory amplification (Chen *et al*, 2024; Dai *et al*, 2021; Lee *et al*, 2023; Mao Zhao & Sun, 2025; Zhang *et al*, 2025), and they position the GSK3β-NF-κB axis as a central node through which FGF10 delivers hepatoprotection and exerts hepatocyte-first control over fibrogenic signaling..

These findings refine the division of labor within hepatic FGF signaling. Endocrine FGFs, such as FGF19 and FGF21, act via FGFR4/FGFR1c with β-Klotho to modulate bile acid and metabolic homeostasis and are in clinical translation for MASLD (Beuers *et al*, 2025; Rose *et al*, 2025; Xu *et al*, 2023). By contrast, paracrine FGFR2b ligands, such as FGF7 and FGF10, have been studied mainly in epithelial regeneration (Mauduit *et al*, 2022; Rice *et al*, 2004), with limited definition in fibrogenesis. Demonstrating that hepatocyte FGFR2 is necessary for the antifibrotic actions of FGF10 assigns an intrinsic epithelial function to the FGFR2 pathway in chronic liver disease and complements the metabolic roles attributed to FGFR4 (Li *et al*., 2024; Lu *et al*, 2019). Our data further indicate that selective reinforcement of FGFR2 under FGF10 delivery coincides with reduced profibrotic transcriptional programs, which is consistent with a receptor-centered mechanism of action in which hepatocytes act as upstream organizers of the fibrotic niche.

An additional nuance is the temporal and compartment-specific regulation of FGF10. In acute or short-term toxicant injury, including prior reports and our early CCl_4_ time-course experiments, hepatic FGF10 and HSC *Fgf10* transcripts are transiently induced, consistent with a compensatory pro-regenerative response. With sustained CCl_4_ exposure and progression to advanced fibrosis, however, both whole-liver FGF10 and HSC *Fgf10* expression follow a biphasic pattern and ultimately decline, a behavior recapitulated by prolonged CCl_4_ treatment in vitro. These kinetics provide a mechanistic basis for reconciling increased FGF10 in acute injury with the stage-dependent FGF10 gradient observed here, whereby transient activation in HSCs and parenchyma gives way to chronic exhaustion or reprogramming of the FGF10-FGFR2 axis in late-stage disease and progressive loss of this endogenous hepatocyte-centered brake.

An additional inference is that temporal, histologic improvement can precede biochemical normalization. In CCl_4_ injury, FGF10 reduced bridging fibrosis, α-SMA, and macrophage infiltration despite limited changes in serum aminotransferases. This pattern is consistent with recompensation (Feng *et al*, 2024), in which structural repair and matrix remodeling advance before full normalization of transient injury markers. Together with reports in acute injury paradigms,(Li *et al*., 2021) these data suggest that FGF10 promotes a conserved pro-survival and pro-regenerative program while dampening inflammation. The observation of regression in collagen burden and fibrosis stage relative to time-matched controls supports the interpretation of partial reversal rather than attenuation alone.

In a metabolic-toxicant context, combining high-fat diet plus CCl_4_, FGF10 retained antifibrotic efficacy and was accompanied by selective metabolic remodeling, including downregulation of *Fasn* and *Srebf2*. Transcriptomic enrichment of lipid and fatty acid pathways after AAV-*Fgf10* or recombinant human FGF10 treatment supports a model in which epithelial protection is coupled to targeted metabolic tuning (Appendix Fig. S16), thereby reducing the inflammatory and profibrogenic burden in steatohepatitis. These findings demonstrate preserved antifibrotic efficacy under metabolic priming and provide evidence of regression beyond a pure toxicant context and suggest that hepatocyte-centered reversal mechanisms can be maintained even in complex metabolic disease.

These results highlight hepatocytes as actionable upstream controllers of the fibrotic niche (Guilliams *et al*, 2022; Yan Cui & Xiang, 2025). Rather than targeting HSCs alone, FGF10-FGFR2-mediated reprogramming of hepatocyte secretory tone reduces TGF-β1 and pro-inflammatory cytokines, imposing a paracrine brake on HSC activation (Filliol *et al*, 2022; Martínez García de la Torre *et al*, 2025; Tao *et al*, 2023). This hepatocyte-first strategy integrates with emerging paradigms of HSCs plasticity, immune-metabolic crosstalk, and epithelial stress responses as co-drivers of fibrosis (Horn & Tacke, 2024; Kaffe *et al*, 2023). We acknowledge that other cell types such as Kupffer cells, cholangiocytes, and endothelial cells may contribute to the response. To strengthen evidence for paracrine control using disease state cells, we envision an ex vivo co-culture that directly pairs hepatocytes and hepatic stellate cells isolated from injured livers with receptor perturbation in hepatocytes in future studies to more precisely map how hepatocyte-intrinsic FGFR2 signaling reprograms the fibrotic ecosystem.

Limitations should be noted. CCl_4_ preferentially models toxicant-driven fibrosis with bridging septa and does not recapitulate all features of cholestatic, immune-dominant, or pure lipotoxic disease. We sought to increase external validity by incorporating dietary models and human tissues, yet formal testing in lipotoxic and immune-skewed settings remains needed. Given FGFR2 expression in cholangiocytes and progenitor compartments, the epithelial-stromal interface warrants deeper resolution by single-cell transcriptomics and lineage tracing. For translation, PK/PD, receptor selectivity, and long-term safety require further definition, particularly with respect to FGFR2-driven epithelial proliferation in chronically injured livers. A trial design that initiates FGF10 after establishment of advanced fibrosis with time-matched comparators would further quantify reversal kinetics and durability and test whether hepatocyte-centered enhancement of FGFR2 signaling can achieve clinically meaningful regression.

In aggregate, chronic injury downregulates FGF10, weakening hepatocyte FGFR2-FRS2α signaling, reducing GSK3β inhibition, and enhancing NF-κB activity. The resulting pro-inflammatory milieu amplifies hepatocyte loss and promotes HSCs activation and ECM deposition. Restoring FGF10/FGFR2 signaling, via recombinant protein, vector-based delivery, or selective FGFR2 agonists, interrupts this cascade, preserves hepatocyte integrity, and attenuates fibrosis. This pathway complements endocrine FGF therapeutics while emphasizing epithelial repair and inflammation control at the level of hepatocyte injury. The requirement for hepatocyte FGFR2 together with histologic regression provides mechanistic and phenotypic support for a hepatocyte-centered fibrosis reversal strategy.

In summary, we identify an intrinsic FGF10-FGFR2 axis that constrains liver fibrogenesis. Across human and murine systems, hepatic FGF10 decreases with stage, and augmenting FGF10 reduces collagen, α-SMA, and inflammatory infiltration, including in metabolically primed disease, and these benefits require hepatocyte FGFR2. Mechanistically, FGF10 restores FGFR2 phosphorylation, engages FRS2α, increases GSK3β (Ser9) phosphorylation, and suppresses NF-κB, thereby lowering inflammatory injury and apoptosis and indirectly limiting HSCs activation. Together, these findings indicate attenuation and partial reversal of established fibrosis under FGF10, which nominates hepatocyte-centered enhancement of FGFR2 signaling, alone or combined with metabolic agents, as a translational path for reprogramming the fibrotic niche and promoting reversal in advanced liver fibrosis.

## METHODS

### Human liver biopsy studies

Liver tissue was obtained from patients with metabolic dysfunction-associated steatotic liver disease (MASLD) at the First Affiliated Hospital of Wenzhou Medical University and stratified by fibrosis stage (FS0-FS3) by blinded histopathologic scoring (FS0, n = 10; FS1, n = 9; FS2, n = 8; FS3, n = 4). Formalin-fixed, paraffin-embedded (FFPE) sections were used for quantitative immunohistochemistry (IHC) to assess FGF10 abundance and localization. Fresh biopsy material (FS0-FS2) was used for RNA extraction and qRT-PCR. The protocol was approved by the institutional ethics committee (approval no. 2016/246); written informed consent was obtained from all participants. No samples were obtained from executed prisoners or institutionalized individuals. Clinical characteristics are summarized in Appendix Table S1.

### Mice and housing

Male C57BL/6J mice (5-8 weeks) were housed under specific pathogen-free conditions (23 ± 2 °C; 60 ± 5% humidity; 12-h light/dark cycle) with ad libitum access to chow and water. Experimental procedures were approved by the Animal Ethics Committee of Wenzhou Medical University (approval no. wydw2022-0691) and conformed to the Guide for the Care and Use of Laboratory Animals. Mice of the same genotype were randomized to treatment groups; investigators were blinded to group assignment during histologic and image analyses unless stated otherwise.

### Diet-induced model

Eight-week-old males received a 60% kcal high-fat diet (HFD) for up to 20 weeks; livers were collected at weeks 5, 16, and 20.

### Carbon tetrachloride (CCl_4_)-induced fibrosis

Eight-week-old males received intraperitoneal CCl_4_ (0.5 mL/kg; 25% in olive oil) three times per week for 6 weeks; controls received vehicle. For time-course analyses, samples were collected at weeks 0, 1, 2, 4, and 6.

### HFD plus CCl_4_ model

Mice were fed HFD for 20 weeks. During the final 8 weeks, CCl_4_ was administered at the above dose/schedule to induce bridging fibrosis.

### Liver-directed FGF10 overexpression

Hepatotropic overexpression used AAV2/8 encoding mouse *Fgf10* under a CAG promoter (AAV-*Fgf10*) or an AAV-GFP control vector. Five-week-old males received 1.2 × 10^11^ vg via lateral tail vein (100 µL). Mice were rested for 3 weeks before fibrosis induction. Vector details and lot numbers are provided in the Key Resources Table (KRT).

### Hepatocyte-Specific Fgfr2 Knockout

*Fgfr2*^flox/flox^ (*Fgfr2^FF^*) mice were crossed with Alb-Cre mice to generate hepatocyte-specific knockout mice (*Fgfr2^LKO^*). PCR-based genotyping confirmed the successful generation of *Fgfr2^LKO^* mice (Appendix Fig. S11). The primer sequences are listed in Appendix Table S2. Knockout efficiency at the protein level in liver was validated by Western blot using anti-FGFR2 antibodies (Appendix Table S3). Age-matched *Fgfr2^LKO^* and *Fgfr2^FF^*males underwent CCl_4_ injury as above.

### Recombinant human FGF10 (rhFGF10) dosing in vivo

During CCl_4_ injury, mice received daily subcutaneous rhFGF10 (1, 2, or 5 mg/kg in 0.9% saline; 5 µL/g) on the dorsal surface; controls received vehicle. Endotoxin content was < 5 EU/mg and bioactivity was 2.8 × 10^5^ AU/mg.

### In vivo optical imaging

To verify liver-restricted transduction, mice received AAV-GFP or saline and were imaged at 0-4 weeks on an IVIS Lumina III under isoflurane anesthesia. Fluorescence was acquired with 480-nm excitation/520-nm emission using identical settings. At week 4, major organs were excised for ex vivo imaging under matched exposure. Radiant efficiency was quantified in Living Image software and rendered with the “YellowHot” colormap using individualized scale limits.

### Serum biochemistry and organ indices

Mice were anesthetized (pentobarbital 100 mg/kg i.p.). Blood from the retro-orbital sinus was allowed to clot, centrifuged (3,000 × g, 30 min, 4 °C), and serum ALT/AST were measured with commercial kits. Body weight was recorded immediately before terminal procedures. After cardiac perfusion with saline, livers were excised, photographed on a standardized background, weighed, and liver-to-body-weight ratio was calculated.

### Histology and morphometry

Liver tissue was fixed in 4% paraformaldehyde, processed to FFPE, and sectioned at 5 µm for hematoxylin and eosin (H&E) and Sirius Red (SR) staining following standard protocols. For lipid staining, liver was embedded in optimal cutting temperature (OCT) compound, snap-frozen, and cryosectioned at 10 µm for Oil Red O. Images were captured with a Nikon ECLIPSE NI microscope and analyzed using Image-Pro Plus software. Fibrosis was quantified by SR-positive area fraction and/or stage scoring as specified in figure legends. Observers were blinded to group.

### Immunohistochemistry (IHC)

FFPE liver sections (5 µm) were deparaffinized, rehydrated, and subjected to heat-induced antigen retrieval in 0.01 M citrate (pH 6.0 buffer) for 20 min. Endogenous peroxidase was blocked with 3% H_2_O_2_ (10 min). After blocking with 5% goat serum in PBS (30 min), sections were incubated overnight at 4°C with primary antibodies. Information on all antibodies (source and dilution) is summarized in KRT and Appendix Table S3. Detection used the ready-to-use SABC-POD (rabbit IgG) kit following the manufacturer’s protocol: incubation with the biotinylated anti-rabbit secondary antibody (30 min, 37°C), then the SABC-POD complex (30 min, 37°C), and visualization with the kit-supplied DAB substrate. Slides were counterstained with hematoxylin, dehydrated, and coverslipped. Imaging was performed on a Nikon ECLIPSE Ni microscope and analyzed with Image-Pro Plus, and quantification used Color Deconvolution (H-DAB) with a fixed threshold across all images in the same batch.

### Immunofluorescence (IF)

After deparaffinization/rehydration and the same antigen retrieval as above, sections were permeabilized with 0.1% Triton X-100 (10 min) and blocked with 5% BSA in PBS (30 min, 37°C). Primary antibodies were applied overnight at 4°C, followed by fluorophore-conjugated secondary antibodies for 1 h at 37°C in the dark. Information on all antibodies (source and dilution) is summarized in KRT and Appendix Table S3. Nuclei were counterstained with DAPI, and slides were mounted in antifade medium. Images were acquired on a Zeiss LSM 980 laser-scanning confocal microscope under identical acquisition settings. Quantification of mean fluorescence intensity was performed using automatic measurement in Zeiss ZEN software with fixed parameters across groups. For cultured cells, fixation was performed with 4% paraformaldehyde (15 min, RT) before permeabilization and blocking as above, and then staining steps were identical to tissue sections.

### TUNEL staining

Apoptotic nuclei were detected using the One-Step terminal deoxynucleotidyl transferase dUTP nick end labeling (TUNEL) Apoptosis Assay Kit. FFPE liver sections (5 µm) were deparaffinized, rehydrated, treated with proteinase K (20 µg/mL, 15 min, RT), and rinsed in PBS. The TUNEL reaction mix was applied for 1 hour at 37°C in the dark, followed by PBS washes. Slides were counterstained with DAPI and mounted in antifade medium. Images were acquired on a Zeiss LSM 980 with identical acquisition settings across groups. The mean fluorescence intensity of TUNEL-positive signals was obtained using automatic measurement in Zeiss ZEN with fixed parameters, and data are presented as relative values.

### RNA sequencing and enrichment analyses

Total RNA was isolated using TRIzol^®^, quantified on a NanoDrop 2000 spectrophotometer, and assessed with an Agilent 5300 Bioanalyzer. Libraries were prepared from RNA meeting the following criteria: OD260/280 between 1.8-2.2, OD260/230 ≥ 2.0, RQN ≥ 6.5, and 28S:18S ≥ 1.0 (Majorbio Bio-pharm Biotechnology Co., Ltd., Shanghai, China). Sequencing was performed on an Illumina NovaSeq X Plus platform (PE150). Reads were aligned to the GRCm39 mouse reference genome using HISAT2 (v2.2.1) and assembled with StringTie (v2.2.0). Differential expression was assessed with DESeq2 (v1.42.0) using Wald tests; genes with |log2FC| ≥ 0.58 and FDR < 0.05 were considered differentially expressed unless stated otherwise. Gene Ontology (GO) enrichment was performed using GOATOOLS (v1.4.4) and pathway enrichment using SciPy-based scripts. All RNA-seq procedures and bioinformatics analyses for the tissue samples were conducted by Shanghai Majorbio Bio-pharm Technology Co., Ltd.

### Isolation and culture of primary mouse hepatocytes and HSCs

Primary mouse hepatocytes and HSCs were isolated from 8-week-old male mice via in situ collagenase perfusion followed by gradient centrifugation as described (Mederacke *et al*, 2015). Briefly, after in situ collagenase perfusion, liver tissue was gently dissociated and filtered. Hepatocytes were collected by low-speed centrifugation, and HSCs were recovered from the supernatant usingdensity-gradient centrifugation. Hepatocytes were cultured in high-glucose DMEM with 10% FBS on collagen-coated plates and identified by albumin immunostaining. HSCs were cultured similarly and identified by desmin, α-SMA, and vitamin A autofluorescence. To minimize passage-related variability, primary hepatocytes were restricted to the first passage, while HSCs were used at passages 1 to 4.

### CCl_4_-induced cell injury and rhFGF10 in vitro

CCl_4_-saturated medium was prepared as follows: 1 mL of CCl_4_ (analytical grade) prefiltered through a 0.22 µm nylon membrane was transferred to an amber serum vial, serum-starved high-glucose DMEM containing 0.5% FBS was added to a final volume of 10 mL, the vial was sealed (butyl rubber stopper plus Parafilm), and incubated at 37°C for 48 h. The upper aqueous supernatant was then collected as the CCl_4_-saturated medium. Primary hepatocytes were exposed to CCl_4_-saturated medium for 6 hour or 24 h as indicated, followed by rhFGF10 (10 or 40 µg/mL). Vehicle controls matched solvent content.

### Annexin V–FITC/PI staining in vitro

Hepatocytes were seeded on collagen-coated glass-bottom dishes and treated as indicated. Without fixation, cells were incubated in Annexin-V binding buffer with Annexin V-FITC and propidium iodide (kit; KRT) for 10 min in the dark, counterstained with DAPI, and imaged immediately on a Zeiss LSM 980. Annexin-V and PI signals were quantified in ZEN using fixed analysis parameters.

### RNA interference

Primary hepatocytes were transfected with si*Fgfr2* or control siRNA using Lipofectamine 3000 for 48 h per manufacturer instructions. Cells were then exposed to CCl_4_-saturated medium ± rhFGF10 for 24 h. Knockdown efficiency was confirmed by immunoblot 72 h post-transfection. siRNA sequences are listed in Appendix Table S4.

### Immunoblotting

Tissue or cell lysates were prepared in lysis buffer with protease/phosphatase inhibitors. Equal amounts of protein (typically 30 µg) was resolved by SDS-PAGE, transferred to PVDF membranes, probed with primary and HRP-conjugated secondary antibodies (KRT and Appendix Table S3), and visualized by chemiluminescence. Band intensities were quantified in Image Lab and normalized to GAPDH.

### Quantitative real-time PCR (qRT-PCR)

Total RNA from liver tissues was extracted using a TRIzol-based kit, reverse-transcribed using Hifair^®^ⅡcDNA synthesis superMix, and subjected to qRT-PCR with SYBR Green master mix on a QuantStudio 5 system. Gene expression was normalized to human *GAPDH* and mouse *Gapdh* using the 2^-ΔΔCt^ method. Primer sequences are provided in Appendix Table S5.

### Statistics

Data are presented as mean ± SEM. Two-group comparisons used two-tailed unpaired Student’s t tests. Multiple groups were analyzed by one-way or two-way ANOVA with appropriate post hoc tests as specified in figure legends. A P value < 0.05 was considered significant. The unit of analysis, biological replicates (n), randomization/blinding, and exact statistical tests are reported in figure legends. All tests were two-sided. Data normality and homoscedasticity were assessed (Shapiro-Wilk, Levene) to select parametric/non-parametric tests. Multiple comparisons were adjusted by BH-FDR where applicable. Effect sizes and 95% CIs are reported when relevant.

## Acknowledgments

This work was supported by the National Natural Science Foundation of China (No. 82400681 to Xuanxin Yang and No.82470066 to Xiaojie Wang), the Xinjiang Safflower Industry Development Fund (to Xiaojie Wang and Xiaokun Li).

## Author contributions

Conceptualization, X.Ya., X.W., X.L.; Formal analysis, B.Yu., H.W., Y.X.; Funding acquisition, X.W., X.Ya., X.L.; Investigation, X.Ya., J.M., Y.Y., Q.D., S.P., Q.W., T.Z., P.Z., Z.J., C.Lu., L.L., X.Yo., Z.Mu.; Methodology, X.Ya., B.Yu., J.M., Y.Y.; Project administration, X.Ya., X.W.; Resources, Y.Y., B.Yu., Q.H., M.Zh., Z.Mu.; Supervision, X.W., X.L., M.Zh.; Validation, X.Ya., B.Yu., J.M., Y.Y.; Visualization, X.Ya., J.M., Y.Y., Y.X.; Writing-original draft, X.Ya., X.W.; Writing-review & editing, X.Ya., X.W., B.Yu., H.W., J.B., Q.H. Equal contribution: X.Ya., B.Yu., J.M., and Y.Y. contributed equally to this work. Initials disambiguation: X.Ya.=Xuanxin Yang; B.Yu.=Bingjie Yu; X.Yo.=Xinyi You; M.Zh.=Minghua Zheng; Z.M.=Zhixiang Mu.

## Disclosure and competing interest statement

The authors declare that they have no conflict of interest.

### The Paper Explained

#### Problem

Advanced liver fibrosis is the main reason why patients with metabolic dysfunction-associated steatotic liver disease (MASLD) develop liver failure and die, yet there is still no approved drug that can reliably reduce this scarring. Fibroblast growth factor 10 (FGF10) helps epithelial repair in other organs, but its role in chronic liver injury is unclear. In particular, it was not known whether an FGF10 pathway in hepatocytes, acting through fibroblast growth factor receptor 2 (FGFR2), can protect the liver and limit fibrosis.

### Results

We found that levels of FGF10 and FGFR2 in the liver fall as fibrosis becomes more severe in patients with MASLD and in mouse models of toxic and metabolic liver injury. When we restored FGF10 specifically in the liver, either with an AAV vector or with recombinant human FGF10, liver inflammation, hepatocyte death, stellate cell activation, and collagen deposition all decreased, and established bridging fibrosis was partly reversed. Single cell and tissue analyses showed that FGF10 is mainly produced by hepatic stellate cells, whereas FGFR2 is expressed in hepatocytes, which supports a paracrine circuit between these cells. Deleting FGFR2 only in hepatocytes largely removed the benefits of FGF10, showing that hepatocyte FGFR2 is required for these antifibrotic and anti-inflammatory effects.

### Impact

Our findings identify a hepatocyte centered FGF10-FGFR2 pathway as a natural protective mechanism against liver scarring that weakens in advanced disease. Strengthening this pathway with FGF10 based treatments or other FGFR2 targeted approaches could offer a new way to reduce inflammation, protect hepatocytes, and promote regression of advanced liver fibrosis, including in metabolically primed steatohepatitis. This hepatocyte focused strategy could complement existing metabolic therapies and broaden the options for patients with MASLD and severe fibrosis.

### Data Availability Section

The data that support the findings of this study are available from the corresponding author upon reasonable request. Raw and processed RNA-seq data will be deposited in GEO upon acceptance; analysis scripts are available upon request. All necessary data that support the findings and conclusions made in this study are included within the article and the supplementary materials.

## Supplemental information

Additional supporting information may be found in the online version of the article at the publisher’s website or from the corresponding author.

